# Deciphering the role of the non-active site ancillary residues in maintaining the activity and substrate specificity of OXA-232 beta-lactamase

**DOI:** 10.64898/2026.05.22.727341

**Authors:** Tejavath Ajith, Bayomi Biju, Diamond Jain, Chiranjit Chowdhury, Anindya Sundar Ghosh

## Abstract

OXA-232, an OXA-48 like carbapenemase stands amongst newly identified beta-lactamases that causes of the extensive of beta-lactam resistance. While active-site residues are well characterised, the contributions of conserved non-active-site residues in exerting enzymatic activity remain unexplored, limiting our understanding about the roles of these residues in the overall OXA-232 function. To address these gaps, the conserved residues S118, V120, L158, and D159 of OXA-232 positioned adjacent to the active-site motifs and within the omega-like loop were substituted with alanine. Substitutions of S118A and D159A rendered the expressing cells susceptible to penicillins, cephalosporins, and carbapenems, whereas the cells harbouring OXA-232_V120A_ and OXA-232_L158A_ proteins exhibited substrate-selective susceptibility changes. Kinetic analysis with purified proteins revealed the reduction in catalytic efficiency of all the mutants compared to wild-type protein. Though the L158A and D159A mutated proteins become deacylation-deficient, the mutations S118A and V120A exhibited selective acylation defects without trapping intermediates. It is evident from circular dichroism spectroscopy and molecular dynamics simulations that OXA-232_S118A_, OXA-232_V120A_, and OXA-232_L158A_ nearly retained their secondary structures and compactness, except for OXA-232_D159A_, which presumably triggered a misfolding leading to destabilisation of the omega-loop. Interestingly, bicarbonate supplementation partially rescued the lost activities in soluble mutants, underscoring the carbamylation dependence. Taken together, these findings establish S118 and D159 as essential for core catalysis and structural integrity, with V120 and L158 modulating substrate-specific turnover and orientation. The current study reappraised the mechanistic insights of OXA-48-like carbapenemases, providing significant resources in rationally designing future therapeutics to combat carbapenem resistance.

---

**MAIN**

Beta-lactam antibiotics are the most widely used antimicrobial agents in medicine, distinguished by a four-membered beta-lactam ring in their molecular structure^1,2^. They act by inactivating penicillin-binding proteins (PBPs), which are essential to cross-link nascent peptidoglycan monomers in both Gram-negative and Gram-positive bacteria^3,4^, hence weakening the bacterial cell wall and ultimately leading to cell lysis. The discovery of penicillin by Alexander Fleming in 1928 and its clinical success during World War II^5^ stimulated the search for new beta-lactams, leading to the discovery of cephalosporins in the late 1940s^6^, carbapenems in the 1980s^7^, and monobactams soon after, each with unique resistance profiles, stability, and clinical applications^7^. Despite their lifesaving properties, the widespread use of beta-lactams promoted bacterial resistance through the synthesis of beta-lactamases, enzymes that hydrolyse the beta-lactam ring and inactivate the drugs^8^. Based on sequence homology, these beta-lactamases are classified into four groups (A-D), with A, C, and D employing a serine-based mechanism and class B metallo-beta-lactamases requiring zinc for catalysis^9,10^. These beta-lactamases have evolved to hydrolyse the carbapenems, which are considered the antibiotics of last resort. These carbapenem-hydrolysing beta-lactamases, commonly known as carbapenemases, are widely distributed in *Enterobacteriaceae* and pose a growing threat to the treatment modalities adapted for antimicrobial-resistant infections in hospitals^11–13^. Among these, OXA-48-like carbapenemases are spreading rapidly across Asia, the Middle East, and Europe, heightening morbidity, mortality, and healthcare costs^14–16^. OXA-232, a variant of class D OXA-48-like carbapenemases, was first detected in 2013 from *K. pneumoniae* and *E. coli* isolates in India, which is now widespread across Asia, Europe, Africa, and the Americas^17^. Carbapenem-encoding genes mostly reside on mobile genetic elements such as plasmids and transposons, consequently promoting their rapid dissemination^18,19^. OXA-232 is encoded on a 6,141 bp ColKP3-type non-conjugative plasmid pkNICU5 isolated from a *K. pneumoniae* strain in China^20^ and is often linked to nosocomial outbreaks, particularly within intensive care units, and frequently coexists with resistance determinants such as 16S rRNA methyltransferases (e.g., rmtF), virulence factors (e.g., rmpA2), and the carbapenemase NDM-5^17,21^. The widespread dissemination of such carbapenemases necessitates the development of new therapeutic regimens, thereby emphasising the need to understand their structural basis of biochemical function in order to address the mechanism towards carbapenem resistance^22^. Structurally, OXA-232 shares substantial sequence identity (92.5%) with OXA-48, but differs only by five substitutions (T104A, N110D, E168Q, S171A, and R214S). Among these, the R214S mutation in the β5-β6 loop of OXA-232 affects its substrate access and hydrolytic water recruitment^23^. Though catalysis depends on the conserved S70-X-X-K73 active-site tetrad and carbamylated lysine (K73)^24^, the non-active-site elements such as the β5-β6 loop and omega-like loop, along with conserved motifs might presumably play crucial roles in substrate recognition, specificity, and stability. Lately, several studies on OXA-232 show reduced susceptibility to clavulanic acid, tazobactam, and sulbactam, which further necessitate characterisation of typically unexplored non-active-site elements^25^. Herein, key residues within the putative omega-like loop, conserved motifs, and β5-β6 loop of OXA-232 are substituted with alanine to investigate their contributions to the enzyme’s functionality. Further, both wild-type and mutant recombinant proteins were purified and characterised for their *in vitro* kinetic efficiency, and thermostability, and to assess impact on the N-carboxylation processes of OXA-232.

## Results

### Conserved non-active-site residues orchestrate OXA-232 catalysis and substrate recognition

OXA-232 shares conserved sequence motifs essential for catalysis with other OXA-48-like family **(Fig S1) (Fig. 1a).** In these class D enzymes, the active site S70-XX-K73 motif drives a two-step hydrolytic process where carbamylated K73 recruits the catalytic water molecule for deacylation **(Fig. 1b)**^26^. The carbamylated K73 is further stabilised by interactions with the residues W157, L158 and D159 of a short alpha helical segment, within the omega-like loops of OXA-23 and OXA-58 **(Fig. 1c)**^27,28^. These conserved hydrophobic and aromatic residues of the omega-like loop enclose the active site and modulate its electrostatic and steric environment for adequate substrate accommodation. The L158 residue mediates hydrophobic contacts that anchor beta-lactam cores and R-groups. At the same time, D159 provides electrostatic stabilisation through salt bridges with β5-β6 loop residues such as R214 **(Fig. 1b)**^29^. The substantial structural similarity of these residues to their counterparts in OXA-23 and OXA-48 highlights their conserved contribution to substrate recognition and hydrolysis. Given that natural variants like OXA-933 (D159N) demonstrate how alterations in residues interacting with the β5-β6 loop can alter carbapenemase behaviour by disrupting key contacts, such as the R214-D159 salt bridge, these positions represent essential sites for functional variation in OXA-48-like enzymes^29^. The S118-X-V120 motif contributes substantially to catalysis, as S118 forms a hydrogen bond with the functional groups of beta-lactam antibiotics, while V120 forms van der Waals contacts with the carbapenem hydroxyethyl side chain to refine active site geometry and interacts with the leucine of omega-like loop to access the water molecule towards the active site during hydrolysis **(Fig. 1b)**^30,31^. Therefore, to assess the impact of these side chains on the hydrolytic performance of OXA-232, alanine substitutions were introduced at S118, V120, L158, and D159, thereby eliminating side-chain effects while retaining the backbone fold **(Fig. 1d).** Further, the wild-type and mutant proteins were purified and evaluated for catalytic efficiency, thermal stability, and effects on N-carboxylation.

**Figure 1:**
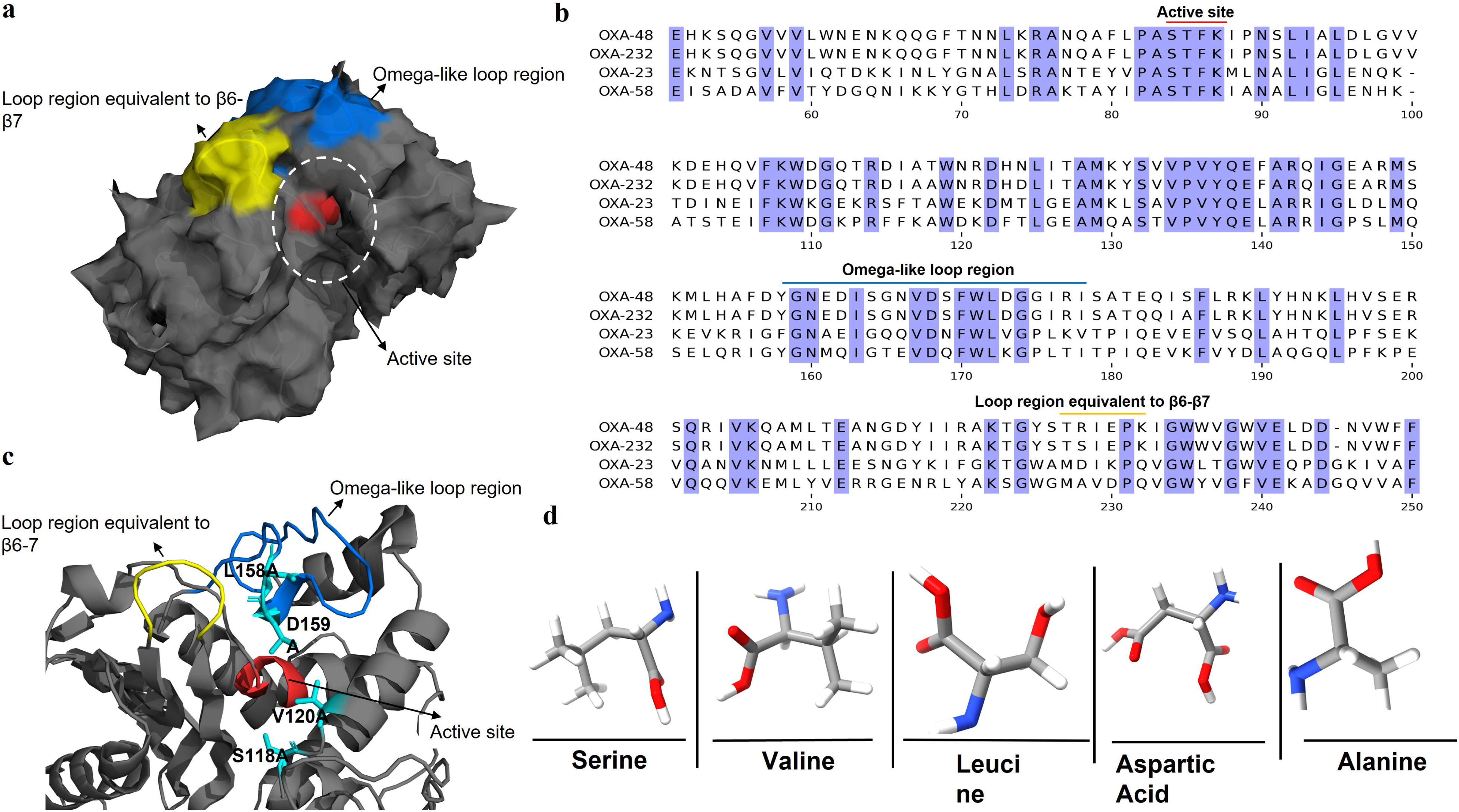
Structural features of OXA-232 guiding mutagenesis. **(a)** Multiple sequence alignment of OXA-232 with OXA-48, OXA-23, and OXA-58, highlighting conserved motifs including the active site, motif II, omega-like loop, and β5-β6 loop regions. **(b)** Detailed view of the active site showing positions of S118, V120, L158, and D159 relative to the omega-like loop, conserved motif II, and carbamylated K73. **(c)** Mutated residues (serine, valine, leucine, aspartic acid) alongside alanine substitutions, illustrating replacement by neutral alanine to eliminate side-chain interactions while preserving backbone conformation.

### Alanine substitutions at S118 and D159 abolished broad-spectrum beta-lactam resistance, while those at V120 and L158 showed substrate-selective susceptibility changes

To evaluate the functional consequences of the S118A, V120A, L158A, and D159A substitutions in OXA-232, we determined the antibiotic susceptibility profiles of *E. coli* AM1OC cells producing the wild-type enzyme and its mutated counterparts against a panel of beta-lactams **(Table 1).** Expression of wild-type OXA-232 conferred substantial resistance, elevating MICs by 16-fold or more for penicillins, cephalosporins, and carbapenems, consistent with its known broad-spectrum hydrolytic capability (Table 1). Conversely, the detectable beta-lactamase activity was abolished in the cells carrying OXA-232_S118A_, making it comparable to wild-type (Table 1). Reduction of resistance indicates underlying essential catalytic role of S118 in the acylation process. Similarly, the cells expressing OXA-232_D159A_ exhibited profound impairment, yielding increased susceptibility towards various beta-lactams, namely, >128-fold for amoxicillin, 8- to 32-fold for other penicillins and cephalosporins, and ∼4-fold for carbapenems, thus confirming the critical contribution of D159 to enzymatic performance and active-site integrity. In contrast, the cells expressing OXA-232_V120A_ retained the resistance, which was comparable to the cells expressing wild-type OXA-232 against penicillins, second- and third-generation cephalosporins (cefuroxime, cefaclor, and cefoperazone). However, it showed marked reductions for first-generation (>16-fold) and carbapenems (imipenem, 8-fold). On the other hand, the cells expressing OXA-232_L158A_ variant also retained resistance to penicillins, while exhibiting moderate reduction in resistance, with a 16- to 32-fold increase in susceptibility to cephalosporins and a 16-fold increase to carbapenems, thus aligning with L158’s hydrophobic contributions to substrate core orientation. These distinct susceptibility patterns emphasise the indispensable roles of S118 and D159 in maintaining broad-spectrum beta-lactamase activity, while revealing the substrate-selective influences of V120 and L158 on OXA-232 function. Building on these *in vivo* observations, we proceeded to purify the wild-type OXA-232 and OXA-232_S118A_, OXA-232_V120A_, OXA-232_L158A_, and OXA-232_D159A_ for detailed biochemical characterisation of their catalytic parameters and structural stability.

**Table 1.** Minimum inhibitory concentrations (MICs) of beta-lactams against *E. coli* expressing wild-type OXA-232 and alanine substitution variants.

| Antibiotics | MIC (µg mL<sup>-1</sup>) |  |  |  |  |  |
| --- | --- | --- | --- | --- | --- | --- |
|  | *E. coli* | OXA-232 and its mutated variants |  |  |  |  |
|  |  | OXA-232 | S118A | V120A | L158A | D159A |
| Penicillins |  |  |  |  |  |  |
| Amoxicillin | 8 | >1024 | 8 | 1024 | 512 | 8 |
| Ampicillin | 8 | >1024 | 8 | 256 | 512 | 128 |
| Piperacillin | 8 | >1024 | 8 | 1024 | 256 | 32 |
| Oxacillin | 16 | >1024 | 16 | 512 | 256 | 128 |
| Cloxacillin | 16 | >1024 | 16 | 256 | 512 | 64 |
| Cephalosporins |  |  |  |  |  |  |
| Cephalothin | 8 | 256 | 8 | 16 | 16 | 32 |
| Cephalexin | 8 | 256 | 8 | 8 | 8 | 8 |
| Cefadroxil | 8 | 256 | 8 | 16 | 8 | 8 |
| Cefuroxime | 16 | >256 | 16 | >256 | 16 | 16 |
| Cefoperazone | 8 | >256 | 8 | >256 | 8 | 16 |
| Cefaclor | 8 | >256 | 8 | >256 | 8 | 32 |
| Carbapenems |  |  |  |  |  |  |
| Imipenem | 0.5 | >32 | 0.5 | 4 | 2 | 8 |

### Alanine substitution in OXA-232 demonstrated substrate-specific catalytic deficiencies

Nitrocefin, a chromogenic cephalosporin, was used to evaluate the effects of alanine substitutions on the *in vitro* catalytic efficiency of OXA-232 proteins. Briefly, the wild-type protein and the S118A, V120A, L158A, and D159A variants were purified to homogeneity using Ni-NTA based immobilised metal ion affinity chromatography, yielding proteins of expected molecular weight of ∼29 kDa with high purity (Fig. 4a). Wild-type OXA-232 exhibited robust turnover with nitrocefin, which moderately enhanced (∼1.2 fold) by NaHCO_3_ supplementation **(Table 2**, **Fig. 2)**. Conversely, the hydrolytic activity was substantially reduced in all variants relative to wild-type OXA-232 (Table 2, Fig. 2). The OXA-232_S118A_ and OXA-232_D159A_ exhibited severe deficits **(Fig. 2a)**, with ∼3.6-fold and ∼29-fold lower catalytic efficiency, respectively, even with NaHCO_3_ supplementation offering minimal restoration (∼1.2-fold at the best) **(Table 2).** At the same time, OXA-232_V120A_ and OXA-232_L158A_ exhibited some intermediate losses in activity (∼4.4- to 5.3-fold), consistent with their more substrate-selective effects.

**Figure 2:**
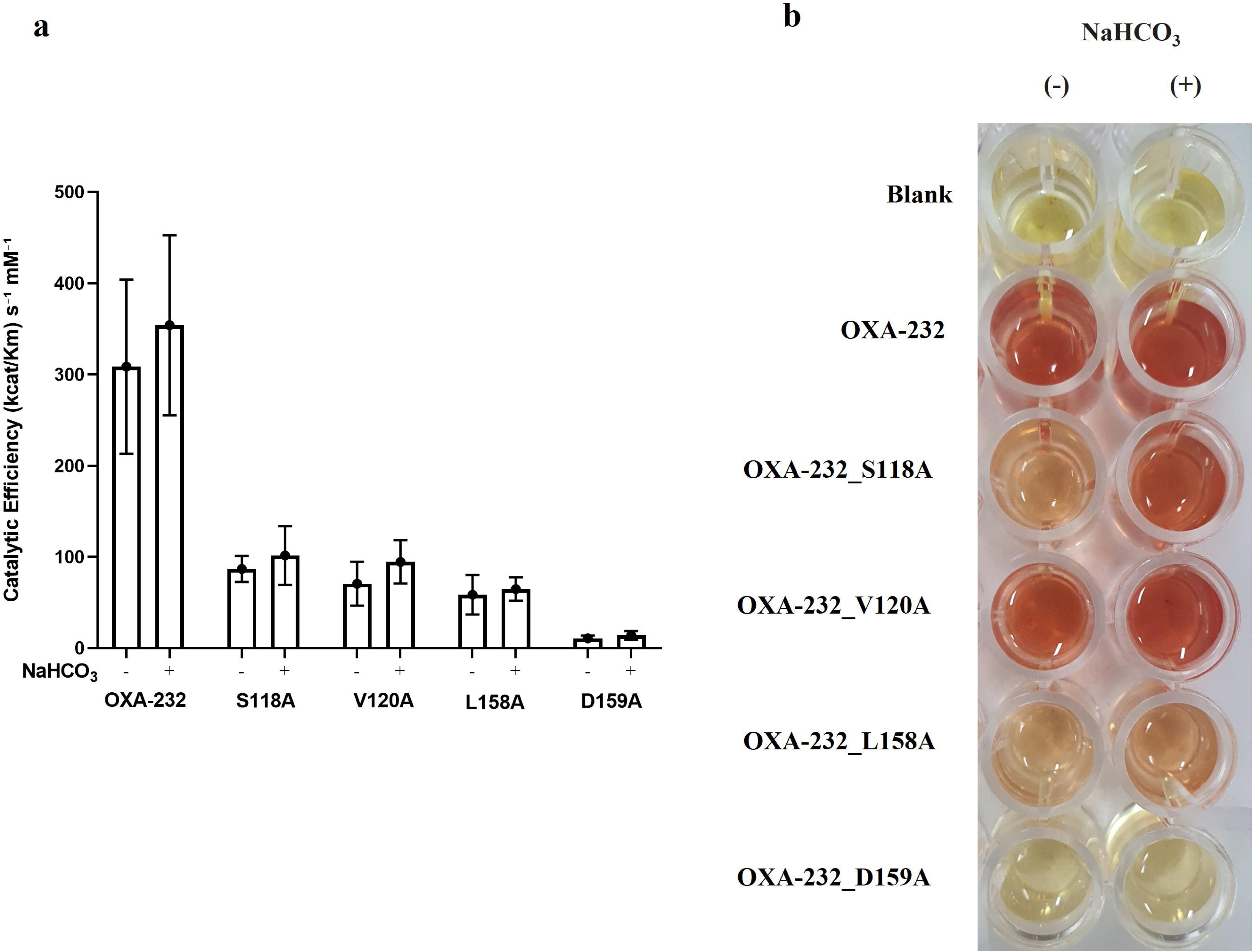
Reduced catalytic efficiency and nitrocefin hydrolysis kinetics of OXA-232 alanine mutants. **(a)** Reduced catalytic efficiency of OXA-232 and mutants measured in the absence and presence of NaHCO_3_, highlighting bicarbonate’s role in modulating class D beta-lactamase activity. **(b)** Visual assay showing nitrocefin hydrolysis colour change by OXA-232 in the presence of NaHCO_3_ compared to buffer blank, confirming active enzyme under bicarbonate conditions.

**Table 2.** Steady-state kinetic parameters of wild-type OXA-232 and alanine mutants toward beta-lactam substrates.

| Antibiotics | Kinetic parameters | OXA-232 | Mutated OXA-232 variants |  |  |  |
| --- | --- | --- | --- | --- | --- | --- |
|  |  |  | S118A | V120A | L158A | D159A |
| Ampicillin | $K_m$ ($\mu\text{M}$) | 345.09 ± 52.68 | ND* | 334.50 ± 47.36 | 310.46 ± 27.59 | ND* |
| | $k_{cat}$ ($\text{s}^{-1}$) | 671.24 ± 83.50 | ND* | 527.09 ± 43.38 | 437.69 ± 22.00 | ND* |
| | $k_{cat}/K_m$ ($\text{s}^{-1} \text{mM}^{-1}$) | 1945.13 ± 371.50 | ND* | 1575.77 ± 258.00 | 1409.79 ± 145.70 | ND* |
| Ampicillin (NaHCO<sub>3</sub>) | $K_m$ ($\mu\text{M}$) | 274.33 ± 60.17 | 137.07 ± 18.66 | 345.09 ± 52.68 | 390.88 ± 64.82 | 48.97 ± 5.00 |
| | $k_{cat}$ ($\text{s}^{-1}$) | 760.46 ± 130.20 | 55.55 ± 4.93 | 671.24 ± 83.50 | 557.14 ± 76.83 | 47.56 ± 2.11 |
| | $k_{cat}/K_m$ ($\text{s}^{-1} \text{mM}^{-1}$) | 2772.05 ± 771.30 | 405.23 ± 65.85 | 1945.13 ± 371.50 | 1425.36 ± 258.00 | 971.09 ± 108.1 |
| Cephalothin | $K_m$ ($\mu\text{M}$) | 217.26 ± 11.18 | ND* | 196.86 ± 34.49 | ND* | ND* |
| | $k_{cat}$ ($\text{s}^{-1}$) | 15.34 ± 0.59 | ND* | 5.30 ± 0.67 | ND* | ND* |
| | $k_{cat}/K_m$ ($\text{s}^{-1} \text{mM}^{-1}$) | 70.7 ± 4.53 | ND* | 26.95 ± 5.83 | ND* | ND* |
| Cephalothin (NaHCO<sub>3</sub>) | $K_m$ ($\mu\text{M}$) | 215.41 ± 21.80 | ND* | 207.05 ± 96.22 | ND* | ND* |
| | $k_{cat}$ ($\text{s}^{-1}$) | 16.18 ± 1.21 | ND* | 11.07 ± 3.77 | ND* | ND* |
| | $k_{cat}/K_m$ ($\text{s}^{-1} \text{mM}^{-1}$) | 75.09 ± 9.45 | ND* | 53.47 ± 30.80 | ND* | ND* |
| Cefadroxil | $K_m$ ($\mu\text{M}$) | 407.23 ± 79.88 | ND* | ND* | ND* | ND* |
| | $k_{cat}$ ($\text{s}^{-1}$) | 6.85 ± 1.13 | ND* | ND* | ND* | ND* |
| | $k_{cat}/K_m$ ($\text{s}^{-1} \text{mM}^{-1}$) | 16.81 ± 4.30 | ND* | ND* | ND* | ND* |
| Cefadroxil (NaHCO<sub>3</sub>) | $K_m$ ($\mu\text{M}$) | 276.96 ± 50.49 | ND* | 270.32 ± 51.95 | ND* | ND* |
| | $k_{cat}$ ($\text{s}^{-1}$) | 4.82 ± 0.69 | ND* | 0.804 ± 0.120 | ND* | ND* |
| | $k_{cat}/K_m$ ($\text{s}^{-1} \text{mM}^{-1}$) | 17.41 ± 4.03 | ND* | 2.976 ± 0.725 | ND* | ND* |
| Cefoperazone | $K_m$ ($\mu\text{M}$) | 199.14 ± 53.44 | ND* | 168.50 ± 23.26 | ND* | ND* |
| | $k_{cat}$ ($\text{s}^{-1}$) | 6.86 ± 1.34 | ND* | 5.14 ± 0.49 | ND* | ND* |
| | $kcat/Km$ ($s^{-1}$<br>$mM^{-1}$) | $34.46 \pm 11.42$ | ND* | $19.84 \pm 3.33$ | ND* | ND* |
| **Cefoperazone**<br>**(NaHCO<sub>3</sub>)** | $Km$ ($\mu M$) | $108.93 \pm 5.04$ | ND* | $123.82 \pm 9.68$ | ND* | ND* |
| | $kcat$ ($s^{-1}$) | $6.26 \pm 0.17$ | ND* | $4.40 \pm 0.22$ | ND* | ND* |
| | $kcat/Km$ ($s^{-1}$<br>$mM^{-1}$) | $57.45 \pm 3.10$ | ND* | $35.5 \pm 3.3$ | ND* | ND* |
| **Imipenem** | $Km$ ($\mu M$) | $55.75 \pm 9.76$ | ND* | ND* | ND* | ND* |
| | $kcat$ ($s^{-1}$) | $2.09 \pm 0.17$ | ND* | ND* | ND* | ND* |
| | $kcat/Km$ ($s^{-1}$<br>$mM^{-1}$) | $37.6 \pm 7.3$ | ND* | ND* | ND* | ND* |
| **Imipenem**<br>**(NaHCO<sub>3</sub>)** | $Km$ ($\mu M$) | $57.29 \pm 6.87$ | ND* | ND* | ND* | ND* |
| | $kcat$ ($s^{-1}$) | $2.302 \pm 0.129$ | ND* | ND* | ND* | ND* |
| | $kcat/Km$ ($s^{-1}$<br>$mM^{-1}$) | $40.18 \pm 5.31$ | ND* | ND* | ND* | ND* |
| **Nitrocefin** | $Km$ ($\mu M$) | $191.31 \pm 48.04$ | $130.13 \pm 18.04$ | $112.97 \pm 32.78$ | $172.77 \pm 52.41$ | $144.95 \pm 33.54$ |
| | $kcat$ ($s^{-1}$) | $59.04 \pm 10.64$ | $11.31 \pm 1.01$ | $7.98 \pm 1.42$ | $10.12 \pm 2.14$ | $1.56 \pm 0.24$ |
| | $kcat/Km$ ($s^{-1}$<br>$mM^{-1}$) | $308.59 \pm 95.38$ | $86.89 \pm 14.31$ | $70.62 \pm 24.02$ | $58.58 \pm 21.68$ | $10.79 \pm 3.00$ |
| **Nitrocefin**<br>**(NaHCO<sub>3</sub>)** | $Km$ ($\mu M$) | $234.75 \pm 65.46$ | $85.91 \pm 23.79$ | $137.59 \pm 28.81$ | $153.10 \pm 25.35$ | $151.07 \pm 42.21$ |
| | $kcat$ ($s^{-1}$) | $74 \pm 15.87$ | $8.73 \pm 1.34$ | $13.01 \pm 1.78$ | $9.93 \pm 1.11$ | $2.11 \pm 0.40$ |
| | $kcat/Km$ ($s^{-1}$<br>$mM^{-1}$) | $354 \pm 98.76$ | $101.58 \pm 32.15$ | $94.58 \pm 23.65$ | $64.83 \pm 12.95$ | $13.98 \pm 4.7$ |

ND\*: Not determined as no hydrolysis detected

Steady-state kinetic parameters (*k*_cat_, *K*_m_, *k_cat_*/*K*_m_) against representative beta-lactams also presented similar trends **(Table 2)**. Wild-type OXA-232 efficiently hydrolysed penicillins, early-generation and third-generation cephalosporins, and carbapenems. The OXA-232_S118A_ exhibited drastic reductions in hydrolytic activity (>20-fold for penicillins, >10-fold for cephalosporins, and >50-fold for imipenem), driven by elevated *K*_m_ values and diminished *k_cat_* values. Intriguingly, Asp159 to Ala substitution completely abolished the activity against most of the substrates (∼28-fold loss for cefoperazone, undetectable activity against cephalothin/cefadroxil and imipenem). On the other hand, an OXA-232_V120A_ variant exhibited null effect for ampicillin and the activity remained similar to that of wild-type. However, the said variant displayed moderate drops for cephalosporins (∼2- to 6-fold) and limited hydrolysis of the carbapenem class (Table 2). The substitution of Leu158 to Ala in OXA-232_L158A_ exhibited severe impairments (>6-fold for penicillins/cephalosporins, complete loss for imipenem) (Table 2). These kinetic deficits was consistent with what we observed for the *in vivo* susceptibility patterns and are in line with the results obtained after nitrocefin hydrolysis.

### Alanine substitutions in the omega-like loop maintain global fold but alter structural resilience

Residues often influence the thermal stability of class D beta-lactamases, particularly those located in or near the active site and within omega-like loop^29,32^. Therefore, we investigated the impact of alanine substitutions at positions S118, V120, L158, and D159 on the secondary structure and thermal stability of OXA-232 using far-UV circular dichroism (CD) spectroscopy. The CD spectra of wild-type OXA-232 exhibited a characteristic alpha-helical profile defined by a positive peak at ∼195 nm and distinct double minima at 208 nm and 222 nm **(Fig. 3a).** Although, both wild-type and mutants represent general spectral signature, overlaying spectral data of the mutant with that of wild-type indicated a large difference between them. Compared to wild-type, there had been notable reduction in molar ellipticity related to the helix components in OXA-232_S118A_. The OXA-232_V120A_, OXA-232_L158A_, and OXA-232_D159A_, on the other hand displayed spectra nearly superimposable on the wild-type, particularly in the presence of sodium bicarbonate (NaHCO_3_), which exerted a normalising effect on the structural signal **(Fig. 3a).** Individual comparisons of each variant in the presence and absence of bicarbonate further illustrate these minor conformational shifts **(Fig. S2a)**.

**Figure 3:**
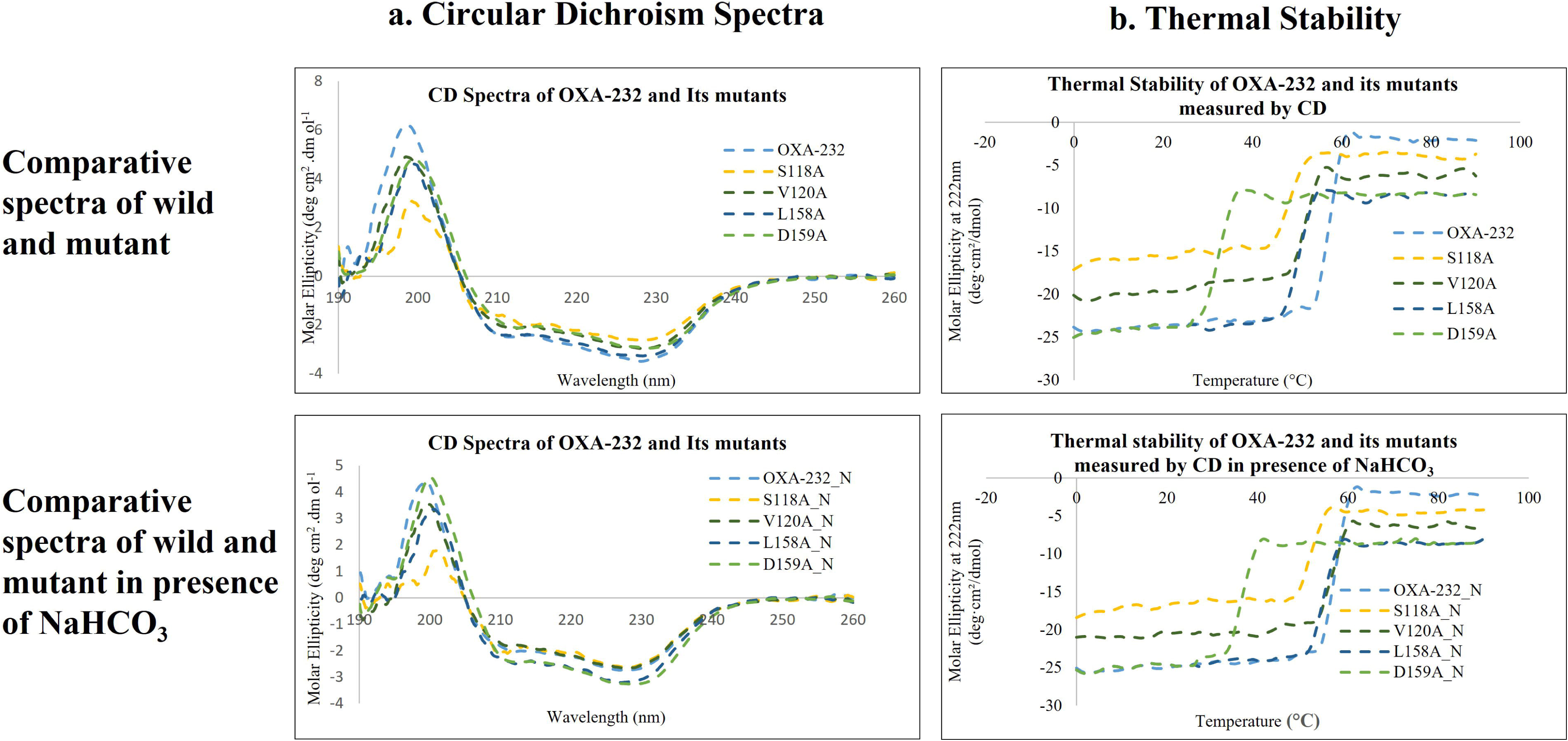
Circular dichroism spectra and thermal stability of OXA-232 variants. **(a)** Circular dichroism (CD) spectra of wild-type OXA-232 and its mutants. **(b)** Thermal denaturation profiles monitored at 222 nm for the same variants. Data were recorded in non-supplemented buffer and in buffer supplemented with 20 mM NaHCO₃.

Temperature-dependent unfolding curves displayed the characteristic sigmoidal profiles indicative of cooperative unfolding **(Fig. 3b).** Individual traces **(Fig. S2b)** indicated that wild-type OXA-232 began unfolding at approximately 50^°^C with a sharp transition and a melting temperature (*T*_m_) of ∼53^°^C. The OXA-232_S118A_, OXA-232_L158A_, and OXA-232_V120A_ showed modest reductions in stability with *T*_m_ values of ∼46^°^C, ∼50^°^C, and ∼52^°^C, respectively. In stark contrast, OXA-232_D159A_ exhibited markedly reduced thermal stability, characterised by a much earlier onset of unfolding (∼30^°^C) and a substantially lower *T_m_* compared to the wild-type and other mutants, both in the absence and presence of NaHCO3 **(Fig. 3b).** Bicarbonate supplementation induced a global stabilising effect, shifting all transitions toward higher temperatures. Specifically, *T*_m_ values increased to approximately 57^°^C for OXA-232, 52^°^C for OXA-232_S118A_, 56^°^C for OXA-232_V120A_, and 55^°^C for OXA-232_L158A_. Notably, while bicarbonate stabilised the OXA-232_D159A_, the unfolding profile remained markedly left-shifted compared to the wild-type, even following supplementation. These comparative unfolding traces confirm that the wild-type remains the most thermally robust variant and bicarbonate consistently induces a rightward shift in stability while preserving close structural alignment between the proteins. These observed changes likely arise from altered local interactions within the active site and the omega-like loop, highlighting the critical role of these regions in maintaining the enzymes’ structural integrity.

### L158A and D159A substitutions render OXA-232 deacylation-deficient with persistent Bocillin-FL acyl-enzyme intermediates

To determine whether reduced hydrolytic activity in OXA-232 mutants stems from impaired acylation or a failure in the subsequent deacylation step, we performed a Bocillin-FL binding assay^33,34^. Purified wild-type and mutant proteins were incubated with the fluorescent penicillin probe. The reactions were quenched at 30 s and 60 s to capture the formation of covalent intermediates by SDS-PAGE and fluorescence imaging **(Fig. 4a).** Wild-type OXA-232 and OXA-232_V120A_ displayed no detectable fluorescent bands at either time point, indicating rapid substrate turnover where the acyl-enzyme intermediate is transient and immediately hydrolysed. Similarly, OXA-232_S118A_ displayed no binding, suggesting that the initial acylation step is severely compromised **(Fig. 4b).** In contrast, OXA-232_L158A_ and OXA-232_D159A_ exhibited intense fluorescent bands that persisted at both time points. This accumulation of the acyl–enzyme intermediate suggests a substantial deacylation defect, causing the enzyme to remain retained in the acyl-enzyme state **(Fig. 4b).** A competitive Bocillin-FL binding assay was subsequently employed to evaluate the stability of these acyl-enzyme adducts using ampicillin and cephalothin as competitors. We hypothesised that if L158A and D159A are truly deacylation-deficient, the active site should remain occupied by the first substrate it encounters, preventing subsequent binding. In the first experimental set **(Fig. 4c, lanes marked ‘L’),** pre-incubation with a molar excess of ampicillin or cephalothin effectively blocked subsequent Bocillin-FL binding, as evidenced by the lack of fluorescence. Conversely, in the second set **(Fig. 4c, lanes marked ‘R’),** where the proteins were allowed to react with Bocillin-FL before the addition of excess antibiotics, the fluorescent probe was not displaced. These results confirm that OXA-232_L158A_ and OXA-232_D159A_ form highly stable, near-irreversible adducts with beta-lactam substrates. The inability of the enzyme to release these molecules proves that L158 and D159 are essential for the deacylation phase of the catalytic cycle.

**Figure 4:**
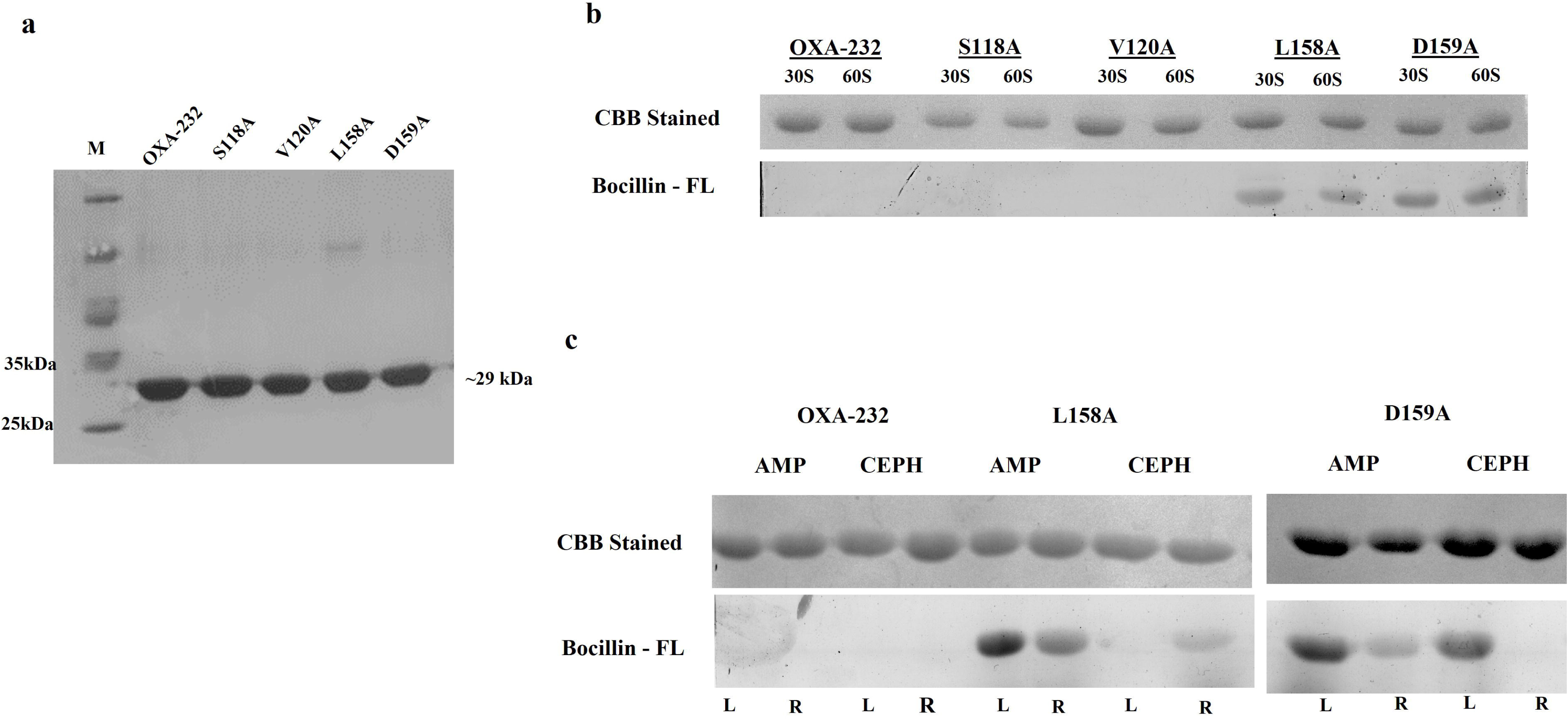
Bocillin-FL binding assay of OXA-232 and alanine mutants. **(a)** Homogeneously purified OXA-232, OXA-232_S118A_, OXA-232_V120A_, OXA-232_L158A_ and OXA-232_D159A_ proteins. **(b)** Bocillin-FL binding assay of purified proteins, revealing persistent acyl-enzyme intermediates for OXA-232_L158A_ and OXA-232_D159A_. **(c)** Competitive Bocillin-FL binding assay of OXA-232_L158A_ and OXA-232_D159A_ with ampicillin and cephalothin. L: antibiotic first, followed by Bocillin-FL; R: Bocillin-FL first, followed by antibiotic.

### Alanine substitutions alter the conformational dynamics and stability of OXA-232 in molecular simulations

To evaluate the impact of alanine substitutions on the structural landscape of OXA-232, 500ns molecular dynamics (MD) simulations were conducted to compare backbone stability, residue flexibility, global compactness, and secondary structure persistence. Wild-type OXA-232 and OXA-232_V120A_ maintained stable backbone trajectories with root-mean-square deviation (RMSD) values ranging between 0.7 and 1.5 Å throughout the production phase **(Fig. 5a).** The OXA-232_S118A_ mutant exhibited noticeable destabilisation with RMSD rising to 1.3 Å after 300 ns. The OXA-232_L158A_ variant showed moderate deviation peaking at 1.5 Å near the 400 ns mark, which correlated with increased localised dynamics within the omega-like loop. Interestingly, the OXA-232_D159A_ displayed a highly rigid backbone profile with an RMSD of ∼0.2-0.3Å despite showing significant deviations in other parameters. Root-mean-square fluctuation (RMSF) analysis further clarified these dynamic shifts. While OXA-232_S118A,_ OXA-232_V120A_, and OXA_232_L158A_ showed mobility comparable to the wild-type at the mutation sites, OXA-232_D159A_ exhibited elevated fluctuations across multiple regions, particularly within the omega-like loop **(Fig. 5b).** Global compactness assessed via Radius of gyration (R_g_) trajectories remained stable for OXA-232_S118A_, OXA-232_V120A_, and OXA-232_L158A_. However, persistent expansion for OXA-232_D159A_ indicates the loss of structural density **(Fig. 5c).** Analysis of secondary structure persistence reflected similar trends. Unlike the wild-type and the other variants, which kept their alpha-helical and beta-sheet frameworks, the OXA-232_V120A_ showed progressive loss of structural integrity **(Fig. 5d).** These findings from the MD simulation mirror the experimental data linking the observed catalytic impairments to the inherent structural fragility of the OXA-232_D159A_ mutant.

**Figure 5:**
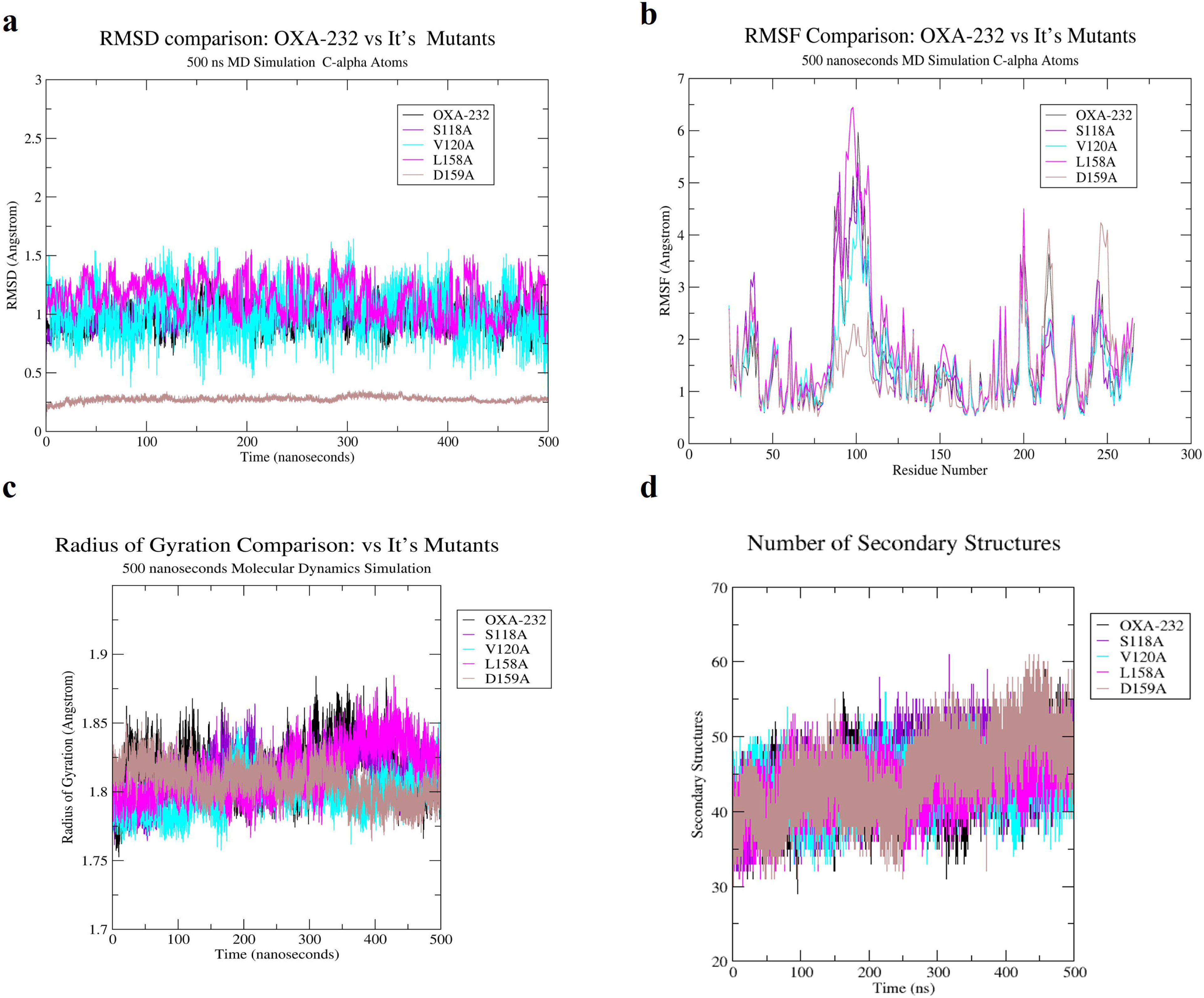
Conformational dynamics of OXA-232 alanine mutants from MD simulations. **(a)** Backbone Root Mean Square Deviation over 500 ns for wild-type OXA-232 and OXA-232_S118A_, OXA-232_V120A_, OXA-232_L158A_ and OXA-232_D159A_. **(b)** Radius of gyration (Rg) trajectories for wild-type and mutants. **(c)** Root Mean Square Fluctuation per residue, indicating flexibility at mutated sites and in the omega-like loop (Y156–I176). **(d)** Secondary Structure persistence profiles over simulation time.

## Discussion

More than eighty years after the Nobel Prize recognised Alexander Fleming’s discovery of penicillin, beta-lactam antibiotics remain foundational to infectious disease treatment. However, their clinical utility is increasingly eroded by the dissemination of carbapenem-hydrolysing class D beta-lactamases (CHDLs). These enzymes are one of the primary drivers of escalating beta-lactam resistance in *A. baumannii* isolates and the *Enterobacterales*^35^. OXA-232, a clinically relevant carbapenemase of the OXA-48-like family (group II) from class D beta-lactamases, exemplifies this challenge^20^. While structural and functional studies have extensively mapped the contributions of active-site residues in OXA-type and other classes of beta-lactamases, the precise roles of conserved non-active-site residues often differ across OXA variants^36–39^. In the present work, we assessed the functional impact of alanine mutagenesis by removing side-chain polarity while preserving main-chain conformation, which revealed distinct molecular determinants for catalysis and stability at conserved motif SXV/I residues S118 and V120 and at omega-like loop residues L158 and D159^38^ within OXA-232, a clinically prominent member of the OXA-48-like family. The residue S118 interacts with functional groups of substrates, specifically acting as a critical proton donor to the beta-lactam nitrogen, similar to its function in OXA-48 and S130 in class A beta-lactamases^28^. Substitution of this residue with alanine (S118A) essentially abolished resistance across all tested beta-lactams due to the disruption of the initial acylation pathway. In contrast, V120 engages in van der Waals contacts with the carbapenem hydroxyethyl side chain^40,41^, acting as a steric brake that stalls deacylation. Interestingly, while the equivalent V167A substitution in OXA-23 was previously reported not to affect susceptibility, our results suggest that the SXV motif in OXA-48-like enzymes plays a more active role in substrate-specific tuning. This was evidenced by the OXA-232_V120A_, which selectively impaired the activity towards first-generation cephalosporins and carbapenems while largely sparing resistance to penicillins and later generation cephalosporins^30^. Within the omega-like loop, L158A is equivalent to L166 in OXA-23, which contributes to hydrolytic function by providing the hydrophobic stabilisation necessary to properly orient the beta-lactam core and R-groups^42^. Substitution at L158 resulted in a significant reduction in MICs against both penicillins and carbapenems. This suggests that the hydrophobic side chain of L158 is critical determinant for substrate recognition and binding performing a role comparable to that of L166 in OXA-23 subfamily^27^. It could likely be due to the loss of van der Waals interactions with the hydroxyethyl moiety of the substrate. Meanwhile D159 establishes salt bridges with β5-β6 loop residues such as R214 to stabilise active-site architecture^31,43^. The significant impairment was observed in OXA-232_D159A_, which resulted a profound loss of resistance to penicillins and carbapenems without eliminating activity, consistent with the effects of D159N/G/K variants in clinical OXA-933 isolates^31^. In these instances, the loss of the electrostatic bridge disrupts the structural integrity required to elicit resistance.

The biophysical characterisation of purified wild-type OXA-232 and its variants through kinetic parameters, Far-UV circular dichroism (CD) spectra and thermal stability analyses reveal a significant reduction in catalytic efficiency that cannot be attributed to global misfolding, as the secondary structure remains preserved^44,45^. Class D beta-lactamases typically exhibit a trade-off between enzymatic activity and thermal stability. In OXA-232, the substitutions at S118 and D159 led to the most pronounced catalytic losses, while L158A produced substrate-selective impairment and V120A showed minimal impact. The stimulatory effect of sodium bicarbonate on all soluble proteins confirms the presence of a functional K73 carbamylation site^46^. However, the differential rescue observed across the mutant variants indicates that these ancillary non-active site residues modulate the efficacy and stability of K73-mediated carbamylation. The formation and maintenance of N-carboxylated K73 in serine beta-lactamases depend heavily on the microenvironment of the surrounding pocket. The substitutions studied here likely disrupt this network by altering the loop confirmations or reducing hydrophobic shielding, which can lead to decarboxylation. Thus, while carboxylation can partially compensate for side-chain losses, optimal hydrolysis remains dependent on a stabilised local scaffold to maintain the functional hydrogen-bonding network. Far-UV CD spectra confirmed that OXA-232_S118A_, OXA-232_V120A_, and OXA-232_L158A_ retain structural profiles nearly identical to wild-type. The markedly destabilised OXA-232_D159A_ maintained its global fold at ambient temperatures yet suffered in structural resilience. The thermal instability of OXA-232_D159A_ results specifically from the disruption of the conserved D159-R214 salt bridge. Unlike the OXA-23 subfamily, which utilises W165 as a hydrophobic anchor for loop orientation, OXA-232 relies on this electrostatic interaction to maintain omega-like loop integrity. The loss of this anchor likely results in a floppy loop conformation that compromises active-site geometry and explains the drastic reduction in catalytic efficiency observed for this mutant. The catalytic defects in S118A and L158A stem from localised perturbations instead of global destabilisation, as evidenced by their largely preserved *T_m_* values. This suggests that while D159 is a structural necessity for maintaining the loop scaffold, S118 and L158 operate as functional necessities where the side chains are required for the chemical steps of acylation and substrate positioning.

The persistent Bocillin-FL labelling observed for OXA-232_L158A_ and OXA-232_D159A_ provides biochemical evidence of a deacylation blockade rooted in the structural destabilisation of the omega-like loop. Unlike the rapid turnover seen in wild-type OXA-232, the accumulation of stable acyl-enzyme intermediates in these mutants confirms that while the active site remains capable of the initial substrate attack, it loses the geometric precision required for the subsequent hydrolytic step. In the case of OXA-232_L158A_, the CASTp-detected volume expansion indicates that the loss of this hydrophobic side chain removes a critical spatial constraint, allowing the beta-lactam core to shift within the pocket. This misalignment effectively shields the ester bond from the catalytic water molecule and resulting in the trapped state, as confirmed by the displacement resistance in our competitive binding assays. Similarly, the deacylation failure in OXA-232_D159A_ underscores the mechanical necessity of the D159-R214 salt bridge, where MD simulations suggest that the loss of this electrostatic anchor increases local loop flexibility. This transformation from a rigid catalytic scaffold into a floppy conformation disrupts the hydrogen-bonding network and the positioning of the deacylation water, thereby preventing the breakdown of the covalent adduct. Whereas, S118 and V120 act as functional tuners within the SXV/I motif, L158 and D159 operate as structural anchors within the omega-like loop that are indispensable for the completion of the catalytic cycle. These findings suggest that the hydrolytic efficiency of OXA-232 is not merely a product of active-site chemistry but is strictly dependent on the long-range structural integrity maintained by these peripheral residues.

The current study accentuates the importance of peripheral residues in maintaining structure and function of OXA-232, thus highlights promising sites for structure-guided inhibitor design against carbapenem-resistant pathogens.

## Conclusion

Alanine substitutions in OXA-232 demonstrate that the peripheral residues S118 and D159 are indispensable for maintaining core catalytic competence and active-site architecture. Notably, the loss of the D159 side chain disrupts a critical electrostatic anchor, leading to significant thermal instability and a total deacylation blockade. In contrast, while V120 and L158 primarily serve to fine-tune substrate specificity and omega-like loop dynamics, the OXA-232_L158A_ variant similarly results in a deacylation deficiency due to active-site volume expansion and substrate misalignment. Despite these localised perturbations, all variants maintain a stable global fold at ambient temperatures, confirming that the observed catalytic failures stem from specific geometric misalignments rather than global misfolding. Collectively, these findings map a network of non-active-site residues that are critical for carbapenemase activity. These interaction networks represent exploitable molecular vulnerabilities for the structure-guided design of next-generation inhibitors aimed at overcoming resistance mediated by OXA-48-like enzymes.

## Materials and Methods

### Bacterial strains, plasmids, and culture conditions

All bacterial strains and plasmids are listed in **Table S1.** Strains were cultured in Lysogeny Broth (LB) broth or agar at 37^°^C for 16–18 hours. For antibiotic susceptibility assays, cation-adjusted Mueller-Hinton (MH) broth was used. All growth media were sourced from HI Media Laboratories (Mumbai, India). Molecular biology reagents were obtained from New England Biolabs (Ipswich, MA, USA). Antibiotics and other chemicals from Sigma-Aldrich (St. Louis, MO, USA) unless specified.

### Cloning, and site-directed mutagenesis

The full-length *bla*_OXA-232_ gene was amplified from *K. pneumoniae* gDNA using specific primers **(Table S2)** and cloned into the arabinose-inducible pBAD18Cm vector under *Nhe*I and *Hind*III restriction sites. Mutations S118A, V120A, L158A, and D159A were created using PFU Turbo polymerase (Agilent Technologies, Santa Clara, CA, USA), following the manufacturer’s protocol, and were confirmed by DNA sequencing (Eurofins Genomics, Bangalore, India). Briefly, the target gene cloned within the plasmid (pBAD18Cm) was amplified using the primers carrying the mutations, followed by *Dpn*I digestion of the native methylated plasmid and then transformed into the maintenance host (XL1Blue). Plasmids were isolated, and the mutations were confirmed by sequencing (Eurofins Genomics, Bangalore, India). Beta-lactamases possess a signal peptide for their periplasmic localisation; therefore, for cytoplasmic protein overexpression, the N-terminal signal peptide of *bla*_OXA-232_ was identified using the SignalP 6.0 server. The primers were designed omitting the nucleotides coding for the signal peptide for cloning the truncated (devoid of signal peptide) gene^48^ into the Isopropyl β-d-1-thiogalactopyranoside (IPTG) inducible pET28a vector at the *Nhe*I and *Hin*dIII sites for purification and subsequent experiments.

### Determination of Minimum Inhibitory Concentrations (MICs)

*E. coli* AM10C strain was transformed with pBAD18Cm constructs **(Table S1)** for MICs by the microbroth dilution method, as described earlier^49,50^. Beta-lactam antibiotics were tested using two-fold serial dilutions, with concentration ranges of 1024–2 µg/mL for penicillins, 512–1 µg/mL for first-generation cephalosporins, 256–0.5 µg/mL for second-generation cephalosporins, and 32–0.06 µg/mL for carbapenems. Each well was made up to 300 μL of MH broth by inoculating with ∼10^5^ cells from an overnight culture, and then incubated at 37^°^C for 16–18 hours. Bacterial growth was measured by OD_600 nm_ using a microplate reader (iMark™ Microplate Absorbance Reader, Bio-Rad, Hercules, CA, USA), and MIC values were interpreted according to Clinical and Laboratory Standards Institute (CLSI) guidelines^51,52^.

### Overexpression and purification of the protein

The *E. coli* BL21(DE3) cells transformed with the pET28a constructs **(Table S1)** were inoculated in 100mL LB media with a specific antibiotic (Kanamycin 50µg/mL) and induced with 0.5 mM IPTG at OD_600 nm_ ∼0.6 at 16^°^C for 16–18 hours for expression. Cells were harvested, resuspended in lysis buffer (10 mM Tris-HCl, 500 mM NaCl, 25 mM imidazole, pH 7.4) and sonicated to lyse the cells. The lysate was then centrifuged, and the supernatant containing 6X-his-tagged protein was collected. Briefly, the supernatant was loaded onto a Ni^2+^-NTA affinity chromatography column (ÄKTA Prime Plus; GE Healthcare, Piscataway, NJ, USA) pre-equilibrated with lysis buffer. Column was washed, and 6xHis-tagged proteins eluted with 25–500 mM imidazole gradient (10 mM Tris-HCl and 500 mM NaCl, pH 7.4). The fractions were subsequently analysed using 12% SDS-PAGE. Homogeneous fractions were pooled and dialysed three times against 10 mM Tris-HCl, 500 mM NaCl (pH 7.4) to remove imidazole.

### Determination of steady-state kinetics

Kinetic parameters (*K_m_, k_cat_, k_cat_*/*K_m_*) of OXA-232 and variants were determined at 25^°^C in 10 mM Tris-HCl and 500 mM NaCl (pH 7.4) buffer, with conditions specified in **Table S3**, to maintain protein stability. Hydrolysis of beta-lactam antibiotics was monitored spectrometrically (BioSpectrometer Kinetics, Eppendorf, Hamburg, Germany). To evaluate the effects of N-carboxylation, which is known to enhance enzyme activity by stabilising the active site, 20 mM sodium bicarbonate was added to the reaction mixture and incubated for 5 minutes before the addition of substrate. Initial reaction velocities (*v_0_*) were calculated from the linear phase of substrate absorbance changes at wavelengths specific to each antibiotic, as detailed in **Table S3.** Kinetic parameters were derived by fitting the initial velocity (*v_0_*) data at various substrate concentrations to the Michaelis-Menten equation using My Curve Fit software. The *k_cat_* was calculated by dividing the maximum velocity (*V_max_*) by the total enzyme concentration. Experiments were performed in triplicate, and the results are expressed as mean ± SD.

### Bocillin-FL binding assays

Bocillin-FL is a fluorescent penicillin derivative, well-established to characterise the penicillin-binding proteins. It has also been used to characterise the deacylation deficiency of beta-lactamases after mutagenesis^53,54^. Therefore, we conducted bocillin binding assay to assess the deacylation deficiency of the mutated OXA-232 proteins. Purified beta-lactamase proteins (2.5 μg per reaction) were mixed with 50 μM Bocillin-FL in a buffered solution of 10 mM Tris-Cl (pH 7.4) containing 500 mM NaCl and incubated at room temperature for either 30 or 60 seconds to enable binding. Reactions were halted by adding SDS-PAGE loading buffer containing sodium dodecyl sulphate (SDS) to denature the proteins and terminate the interaction. 12% SDS-PAGE, then analysed the protein-Bocillin-FL complexes^33,34^. Fluorescent signals from Bocillin-FL bound to the beta-lactamases measure the binding efficiency of beta-lactamases. Typhoon FLA 7000 scanner (GE Healthcare Biosciences, Piscataway, NJ, USA) with appropriate excitation and emission settings for fluorescence detection was used to evaluate this binding efficiency qualitatively. A competitive Bocillin FL experiment was conducted to examine further the impact of mutations on OXA-232 activity against beta-lactam antibiotics. The enzyme was initially incubated with a high concentration of antibiotic for 1 minute, followed by the introduction of Bocillin FL. On the flip side, with simultaneous reactions, Bocillin FL was initially added and incubated for one minute prior to the introduction of the antibiotic. The reactions were subsequently terminated using SDS PAGE loading dye, resolved on 12% SDS PAGE, and scanned with the Typhoon FLA 7000. Any decrease in Bocillin FL fluorescence in the presence of a specific beta-lactam was interpreted as competitive occupancy of the active site, thus offering a relative assessment of antibiotic affinity and the mutation’s effect on binding and catalytic activity.

### Secondary structure and thermal stability analysis using Circular Dichroism (CD) spectroscopy

Far-UV circular dichroism spectroscopy was employed to characterise both the secondary structure and thermal stability of the purified beta-lactamase^25,27^. Proteins were dialysed against 10 mM Tris-HCl, pH 7.4, containing 150 mM NaCl, and in parallel in the same buffer supplemented with 20 mM sodium bicarbonate to assess bicarbonate-dependent effects on conformation and unfolding. Spectra were recorded on a JASCO J-1500 spectropolarimeter using a 0.1 cm quartz cuvette over a wavelength range of 190–260 nm, with matched buffer baselines subtracted from all measurements. Raw ellipticity converted to molar ellipticity [θ] (deg cm² dmol⁻¹) using the standard relationship [θ] = 100 θ / (c l n), where c is the protein concentration in mol L⁻¹, l is the path length in cm. N is the number of residues, allowing direct comparison between spectra. For thermal denaturation experiments, the CD signal at 222 nm, which primarily reports on α-helical content, was monitored. At the same time, the temperature was increased from 0^°^C to 90^°^C at a constant rate of 1^°^C per minute, with data collected at every 0.5^°^C intervals in both buffer systems. Ellipticity versus temperature traces were smoothed using an 11-point Savitzky–Golay filter with a third-order polynomial to reduce noise while preserving cooperative unfolding behaviour, and apparent melting temperatures were calculated from the minimum of the first derivative of ellipticity with respect to temperature, *T*_m_ = arg min(−dθ/dT). All measurements were performed at least in triplicate, yielding *T*_m_ values with standard deviations of less than 0.5^°^C. The processed data were then exported to spreadsheet software for graphical presentation and further analysis.

### Structure modelling and protein dynamic simulations

Multiple sequence alignment was performed using Clustal Omega to analyse sequence conservation and identify functional motifs across the study proteins. The crystal structure template (PDB ID: 5HFO) was retrieved from the RCSB Protein Data Bank and used for all structural modelling and analysis. Structure alignments and inspections were carried out using PyMOL (open source), which was also used for generating all molecular visualisations and publication-quality images. Homology models were constructed using MODELLER^55^. With the 5HFO template, all models were processed with CHARMM-GUI^56^ to assign appropriate protonation states and hydrogen atoms, applying the CHARMM36 force field (2022 release) for system parameterisation. Prepared systems were solvated using the TIP3P water model, neutralised with ions, and subjected to energy minimisation and equilibration under both NVT and NPT ensembles. Production molecular dynamics simulations were conducted using GROMACS^57^ 2022 for 500 nanoseconds under periodic boundary conditions, with long-range electrostatics handled by the Particle Mesh Ewald method and covalent bond constraints enforced by the LINCS algorithm. Trajectories were saved every 250 picoseconds and analysed using standard GROMACS utilities for root mean square deviation (RMSD), root mean square fluctuation (RMSF), radius of gyration, and principal component analysis (PCA). All subsequent figures were prepared using PyMOL (open-source). All computations were performed using the PARAM-SHAKTI supercomputing facility at IIT Kharagpur.

## ASSOCIATED CONTENT

### Supporting Information

**Figure S1 Maximum Likelihood phylogenetic tree of representative beta-lactamases**. OXA-232 (highlighted) clusters strictly within the Class D OXA-48 family, distinct from Class A, B, and C enzymes. Shaded area indicates the OXA-48-like clade. Branch lengths represent amino acid substitutions per site.

**Figure S2 Circular dichroism spectra and thermal stability of OXA-232 alanine mutants. (a)** Circular dichroism spectra of OXA-232 wild-type and mutants in the absence and presence of NaHCO₃. **(b)** Thermal stability profiles measured by CD at 222 nm for the corresponding proteins, comparing conditions without and with NaHCO_3_ supplementation.

**Table S1.** Bacterial strains and plasmids used in this study

**Table S2.** Oligonucleotide primers used for mutagenesis and cloning

**Table S3.** Experimental conditions for steady-state kinetic parameter determination

**Table S4.** Surface and volume analysis of OXA-232 variants using CASTp

**Methodology:** In silico studies: The surface area and volume were predicted using CASTp server (Tian, et al. 2018: W363-W7).

## AUTHOR CONTRIBUTIONS

T.A. was involved in data curation, experimentation, formal analysis, visualization, and preparation of the original draft of the manuscript. B.B. contributed to experimentation, formal analysis, and validation of the study. D.J. assisted in the initial stages of experimentation and study execution. C.C. contributed to validation and review and editing of the manuscript. A.S.G. was involved in conceptualisation, supervision, resource management, funding acquisition, project administration, validation, and final review and editing of the manuscript. All authors have read and approved the final version of the manuscript.

## Notes

The authors declare no competing financial interest

## ABBREVIATIONS

K_m_: Michaelis-Menten constant
kDa: kilo Dalton
k_cat_: turnover number
CD: circular dichroism
UV: ultraviolet
CLSI: clinical laboratory standard institute

## ACKNOWLEDGMENTS

The authors sincerely acknowledge the Central Research Facility, Indian Institute of Technology Kharagpur, for providing the Circular Dichroism facility, Dr. Amitabha Bhattacharjee for providing the clinical isolate harbouring *bla*_OXA-232_, and the Param Shakti Supercomputer facility at Indian Institute of Technology Kharagpur for computational support. The authors also acknowledge the use of AI-assisted tools for language refinement and grammatical editing of the manuscript, while taking full responsibility for the scientific content of the work.

