## supplementary file for "Deciphering the role of the non-active site ancillary residues in maintaining the activity and substrate specificity of OXA-232 beta-lactamase"

**Table S1. Bacterial strains and plasmids used in this study**

| Strain/Plasmid | Genotype/Feature | Reference |
|---|---|---|
| ***Strains*** | | |
| *E. coli* XL1B | *RecAendAhsdRsupEthirecAgyrArelAlac* | (Stratagene, LaJolla, Calif.) |
| *E. coli* BL21(DE3) | *F- ompThsdS (r – B m- B) galdcmλ(DE3)* | (Stratagene, West Cedar Creek, TX) |
| *E. coli* CS109 | *W1485 rpoS rph* | C. Schnaitman |
| *E. coli AM1OC* | *2443 ΔampC* | Lab stock |
| ***Plasmids*** | | |
| pBAD18cm | Expression vector with arabinose inducible promoter | (Guzman, et al. 1995: 4121-30) |
| pET28a (+) | T7 promoter based IPTG inducible vector with 5' end 6X-his tag | Novagen, Madison, WI |
| pDOXA232 | $bla_{\text{OXA-232}}$ cloned in pBAD18Cm | This Study |
| pDOXA232S118A | pBAD18Cm bearing $bla_{\text{OXA-232\_S118A}}$ | This Study |
| pDOXA232V120A | pBAD18m bearing $bla_{\text{OXA-232\_V120A}}$ | This Study |
| pDOXA232L158A | pBAD18cm bearing $bla_{\text{OXA-232\_L158A}}$ | This Study |
| pDOXA232D159A | pBAD18cm bearing $bla_{\text{OXA-232\_D159A}}$ | This Study |
| pTOXA232 | Truncated $bla_{\text{OXA-23}}$ cloned in pET28a (+) | This Study |
| pTOXA232S118A | Truncated $bla_{\text{OXA-232\_S118A cloned}}$ in pET28a (+) | This Study |
| pTOXA232V120A | Truncated $bla_{\text{OXA-232\_V120A cloned}}$ in pET28a (+) | This Study |
| pTOXA232L158A | Truncated $bla_{\text{OXA-232\_L158A cloned}}$ in pET28a (+) | This Study |
| pTOXA232D159A | Truncated $bla_{\text{OXA-232\_D159A cloned}}$ in pET28a (+) | This Study |

### Table S2. Oligonucleotide primers used for mutagenesis and cloning

| Primer | Primer sequence (5' – 3') | Product Details |
|---|---|---|
| FP_OXA-232 | CTCTCTGCTAGCAGGAGGATATATATATGCGTGTATTAGCCTTATCGGC | Full length |
| RP_OXA-232 | CTCTCTAAGCTTCTAGGGAATAATTTTTTCCTGTTTGAGC | |
| FP_sOXA-232 | CTCTCTGCTAGCATGAAGGAATGGCAAGAAAACAAAAGTTGGAATGCTC | Truncated |
| FP_S118A | CGCGATGAAGTACGCAGTTGTGCCTGTTT | Mutants |
| RP_S118A | AAACAGGCACAACTGCGTACTTCATCGCG | |
| FP_V120A | GAAGTACTCAGTTGCGCCTGTTTATCAAG | |
| RP_V120A | CTTGATAAACAGGCGCAACTGAGTACTTC | |
| FP_L158A | AGACAGTTTTTGGGCCGATGGTGGTATTC | |
| RP_L158A | GAATACCACCATCGGCCCAAAAACTGTCT | |
| FP_D159A | CAGTTTTTGGCTCGCTGGTGGTATTCGCA | |
| RP_D159A | TGCGAATACCACCAGCGAGCCAAAAACTG | |

**Table S3. Experimental conditions for steady-state kinetic parameter determination**

| Antibiotic | Concentration range (μM) | Δε ($M^{-1}cm^{-1}$) | λ (nm) | Enzyme (nM) |
|---|---|---|---|---|
| Ampicillin | 50–250 | -820 | 235 | 10-100 |
| Cephalothin | 20-100 | -6,500 | 260 | 10-100 |
| Cefadroxil | 20-100 | -9970 | 260 | 10-100 |
| Cefoperazone | 20-100 | -8460 | 260 | 10-100 |
| Imipenem | 10-50 | -9,000 | 300 | 10-100 |
| Nitrocefin | 20-100 | 15000 | 490 | 10-100 |

**Table S4. Surface and volume analysis of OXA-232 variants using CASTp**

| Protein | Surface area (A$^{O2}$) | Surface volume (A$^{O3}$) |
|---|---|---|
| OXA-232 | 175.955 | 175.716 |
| OXA-232_S118A | 174.710 | 156.347 |
| OXA-232_V120A | 187.161 | 162.958 |
| OXA-232_L158A | 204.601 | 173.310 |
| OXA-232_D159A | 174.409 | 153.321 |

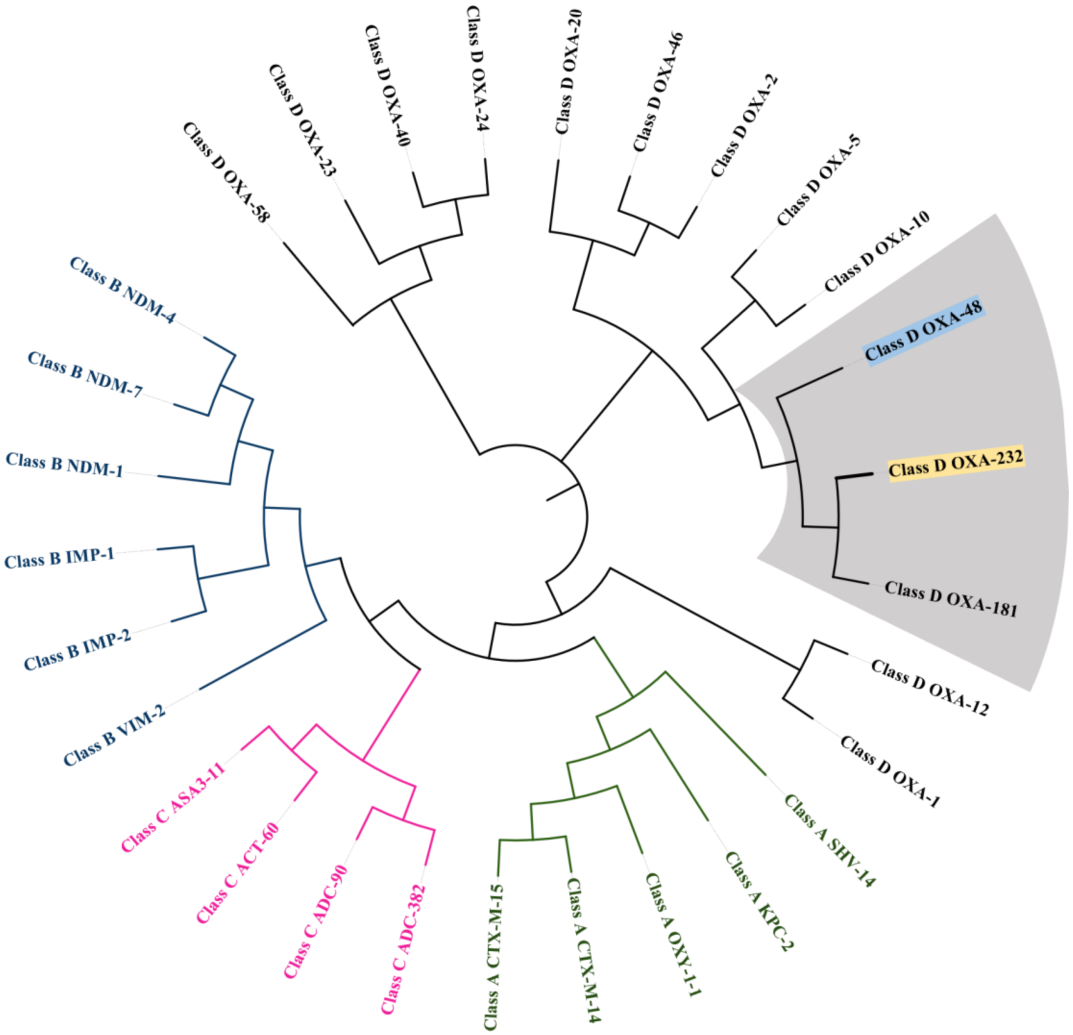


**Figure S1 Maximum Likelihood phylogenetic tree of representative beta-lactamases.** OXA-232 (highlighted) clusters strictly within the Class D OXA-48 family, distinct from Class A, B, and C enzymes. Shaded area indicates the OXA-48-like clade. Branch lengths represent amino acid substitutions per site.

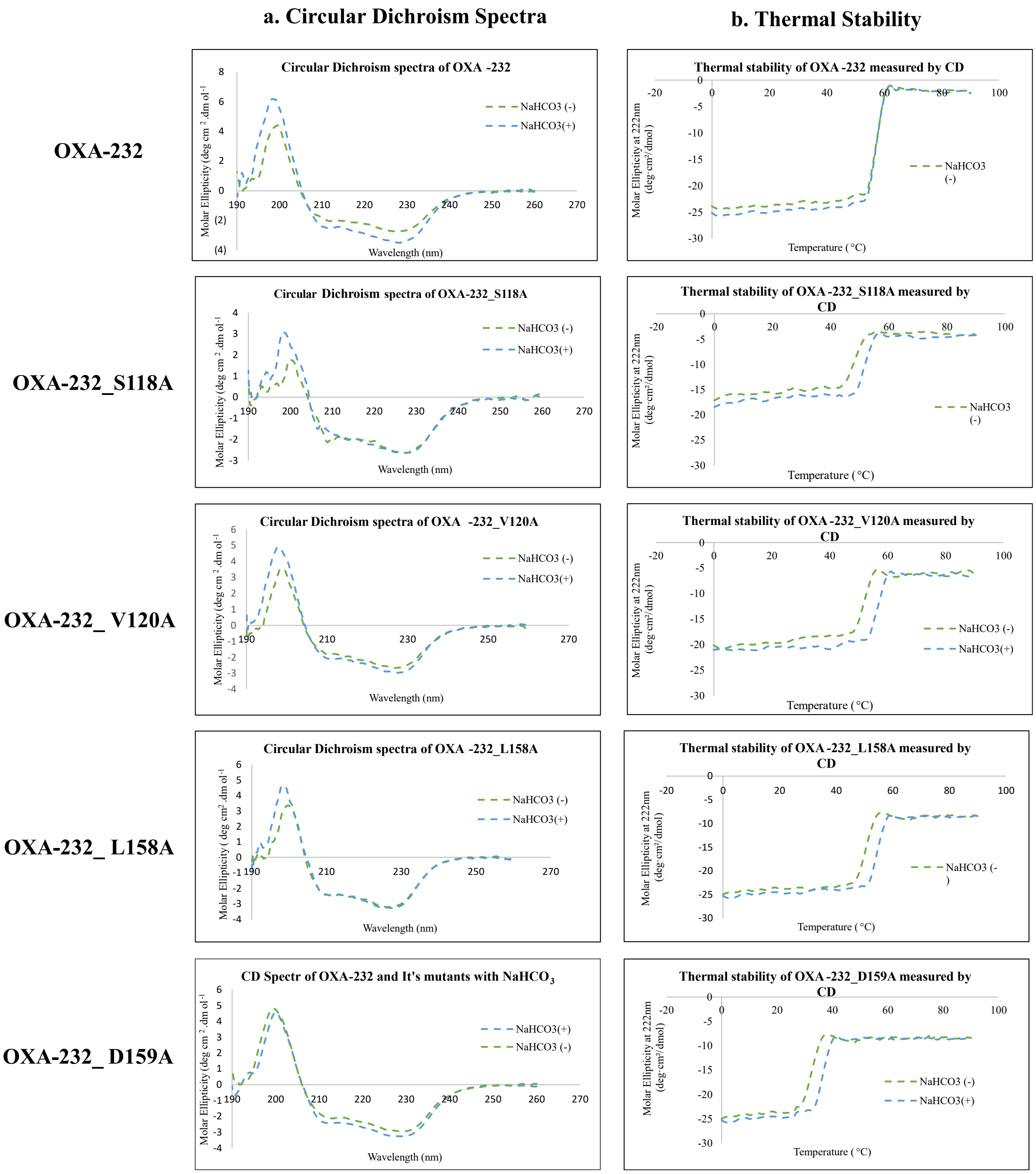


**Figure S2 Circular dichroism spectra and thermal stability of OXA-232 alanine mutants. (a)** Circular dichroism spectra of OXA-232 wild-type and mutants in the absence and presence of $NaHCO_3$. **(b)** Thermal stability profiles measured by CD at 222 nm for the corresponding proteins, comparing conditions without and with $NaHCO_3$ supplementation.

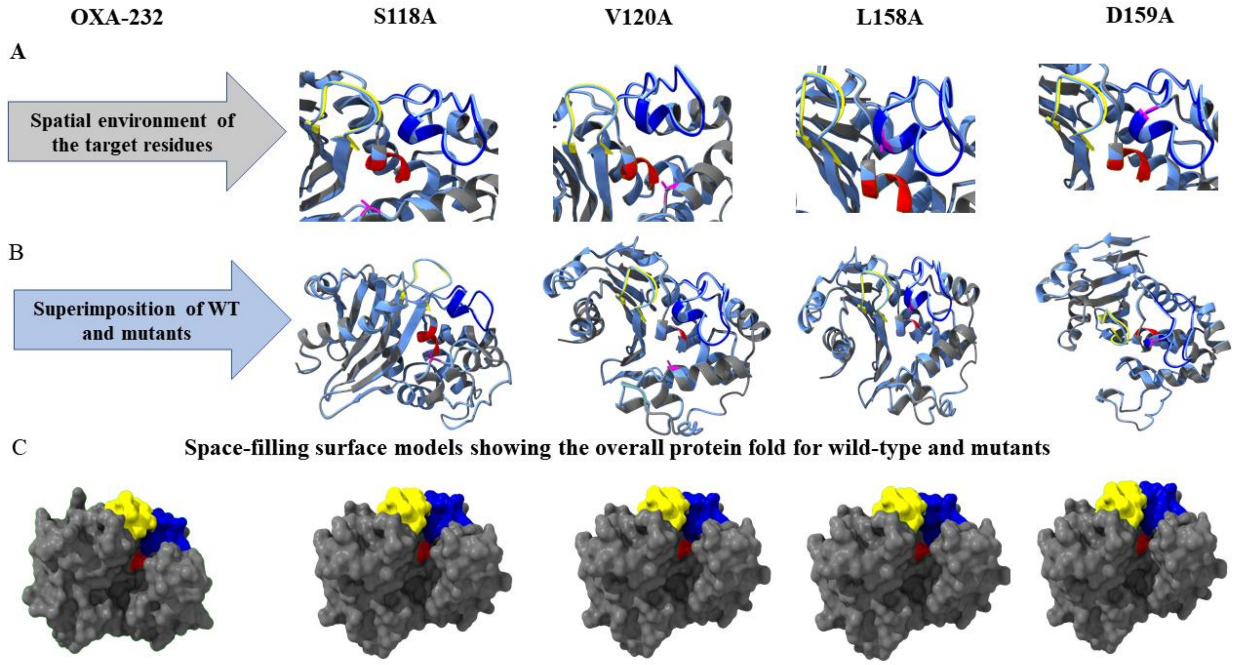


**Figure S 3 Supplementary Figure S3 Structural analysis of OXA-232 mutants**. **(A)** Mutation landscape depicting the spatial location of the mutated residues (pink) relative to the active site (red), Omega-like loop region(blue), and Loop region equivalent to β6-β7 (yellow). **(B)** Structural superimposition of wild-type (gray) and mutant (colored) protein backbones used to visualise localized conformational shifts. **(C)** Space-filling surface models showing the overall protein fold for wild-type and each mutant, illustrating the preservation of the global structural architecture

**Methodology:**

***In silico* studies:** The surface area and volume were predicted using CASTp server(Tian, et al. 2018: W363-W7).
